# Antisense oligonucleotide therapy rescues aggresome formation in a novel Spinocerebellar Ataxia type 3 human embryonic stem cell line

**DOI:** 10.1101/681650

**Authors:** Lauren R. Moore, Laura Keller, David D. Bushart, Rodrigo Delatorre, Duojia Li, Hayley S. McLoughlin, Maria do Carmo Costa, Vikram G. Shakkottai, Gary D. Smith, Henry L. Paulson

## Abstract

Spinocerebellar ataxia type 3 (SCA3) is a fatal, late-onset neurodegenerative disorder characterized by selective neuropathology in the brainstem, cerebellum, spinal cord, and substantia nigra. Here, we characterize the first NIH-approved human embryonic stem cell (hESC) line derived from an embryo harboring the SCA3 mutation. Referred here as SCA3-hESC, this line is heterozygous for the mutant polyglutamine-encoding *CAG* repeat expansion in the *ATXN3* gene within the pathogenic repeat range for SCA3. We observed relevant molecular hallmarks of the human disease at all differentiation stages from stem cells to cortical neurons, including robust ATXN3 aggregation and altered expression of key components of the protein quality control machinery. Finally, antisense oligonucleotide-mediated reduction of ATXN3 prevented the formation of p62-positive aggresomes in SCA3-hESCs. The SCA3-hESC line offers a unique and highly relevant human disease model that holds strong potential to advance understanding of SCA3 disease mechanisms and facilitate the evaluation of possible SCA3 therapies.

**Highlights:**

- Generated first NIH-approved SCA3 human embryonic stem cell line (SCA3-hESC)
- SCA3–hESC exhibit robust ATXN3 aggregation pathology and form p62+ aggresomes
- Anti-ATXN3 antisense oligonucleotides rescue aggresome formation in SCA3-hESC
- Derived SCA3 neurons form aggregates and exhibit impaired protein quality control

## 1. Introduction

Spinocerebellar ataxia type 3 (SCA3) is a late-onset neurological disorder and the most common autosomal dominant ataxia worldwide^1^. One of nine polyglutamine (polyQ) diseases, SCA3 is caused by a heterozygous *CAG* trinucleotide repeat expansion that produces an abnormally long, aggregate-prone polyQ sequence in the encoded disease protein^2^. In SCA3, this polyQ expansion occurs in ATXN3, a deubiquitinase with wide-ranging functions in the ubiquitin proteasome system, macroautophagy, DNA damage repair, and transcriptional regulation^1,3^. Despite ubiquitous expression, polyQ-expanded ATXN3 inflicts neuronal dysfunction and loss in discrete brain nuclei spanning the brainstem, cerebellum, spinal cord, substantia nigra, diencephalon, and striatum through a presumed dominant toxic gain-of-function mechanism^4,5^. There remains limited understanding of the pathogenic cascade leading to neurodegeneration, particularly in prodromal SCA3 stages, as well as a lack of well-supported hypotheses for tissue-specific vulnerabilities, and no approved treatments to slow or stop progression of this fatal disease^1,2^.

Disease-specific human pluripotent stem cell (hPSC) lines, including patient-derived induced pluripotent stem cell (iPSC) lines and human embryonic stem cell (hESC) lines derived from donated disease embryos, are proving to be increasingly important model systems for the study of neurodegenerative diseases^6,7^. Disease-specific hPSC lines enable the study of disease processes in human disease-vulnerable differentiated cellular populations that express endogenous levels of pathogenic genes. In the past decade, several hPSC lines carrying the SCA3 mutation have been developed, including several SCA3 patient-derived iPSC lines^8–18^ and one hESC line harboring the SCA3 mutation^19^. This SCA3 hESC line, however, is not approved for research use in the United States. While the production of these lines represents progress towards improved human disease model systems, few or in some cases conflicting studies have been performed to verify that these developed SCA3 hPSC lines replicate well-established SCA3 molecular phenotypes. Identification of quantifiable, disease-dependent molecular phenotypes is arguably needed in order to use such SCA3 hPSC lines to investigate disease mechanisms or in preclinical testing of potential therapeutic agents for SCA3.

Here we report the validation and characterization of the first SCA3 disease-specific hESC line added to the NIH registry of approved hESC lines, designated UM134-1 on the NIH registry but referenced here as SCA3-hESC. In addition to confirming stem cell-like properties of undifferentiated SCA3-hESC, we evaluate well-established molecular phenotypes of SCA3 in undifferentiated stem cells and in differentiated neural progenitor cells and forebrain cortical neurons. We also demonstrate the potential for SCA3-hESC to serve as a disease model to facilitate preclinical drug development by assessing whether molecular phenotypes are rescued following treatment with an anti-*ATXN3* antisense oligonucleotide (ASO) recently validated in preclinical SCA3 transgenic mouse studies^20,21^. Together the findings support the SCA3-hESC line as an important new biological reagent for the SCA3 field, and establish its potential to improve human SCA3 disease modeling and preclinical drug assessment.

## 2. Materials and Methods

### 2.1 Key Resource Table

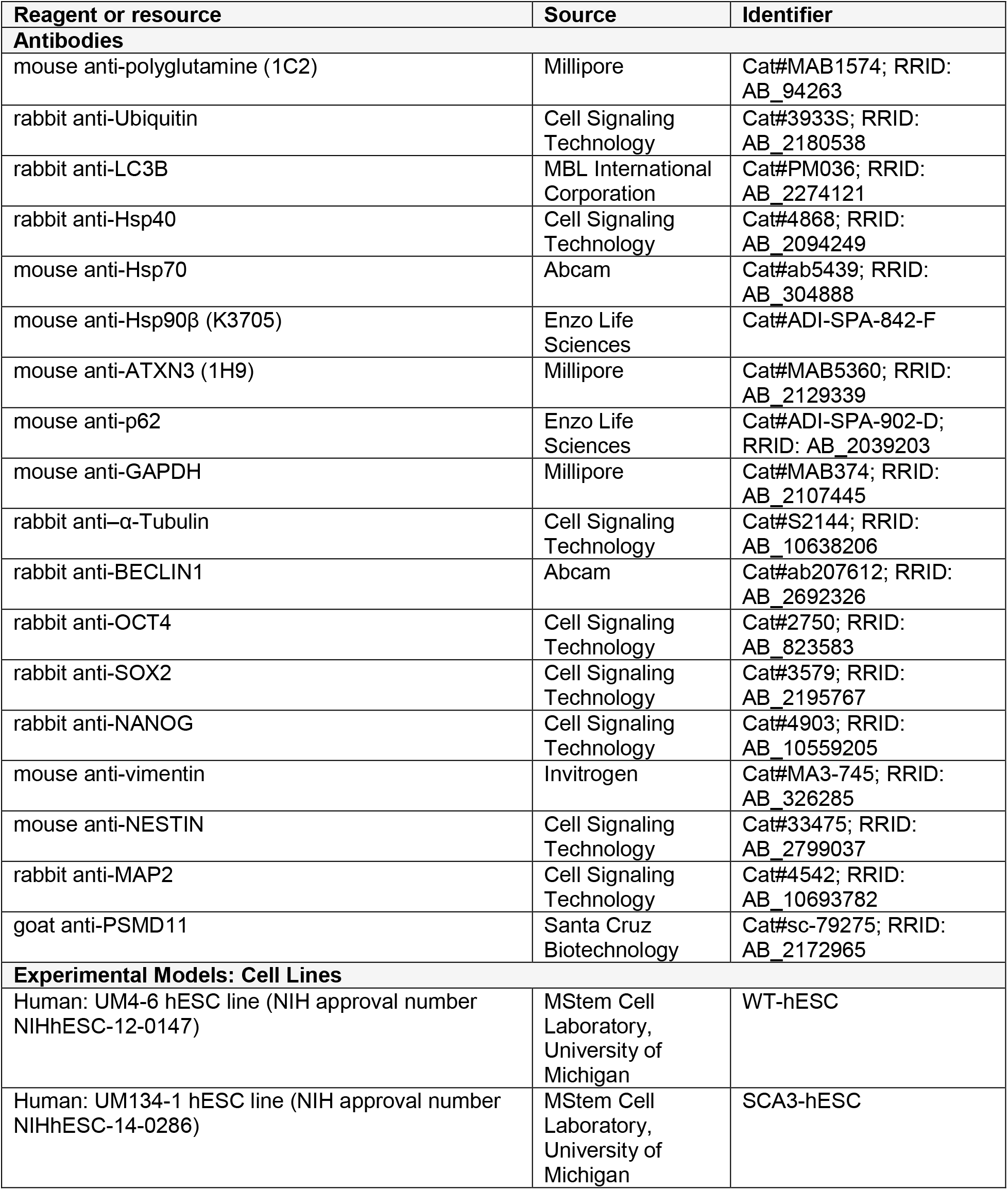
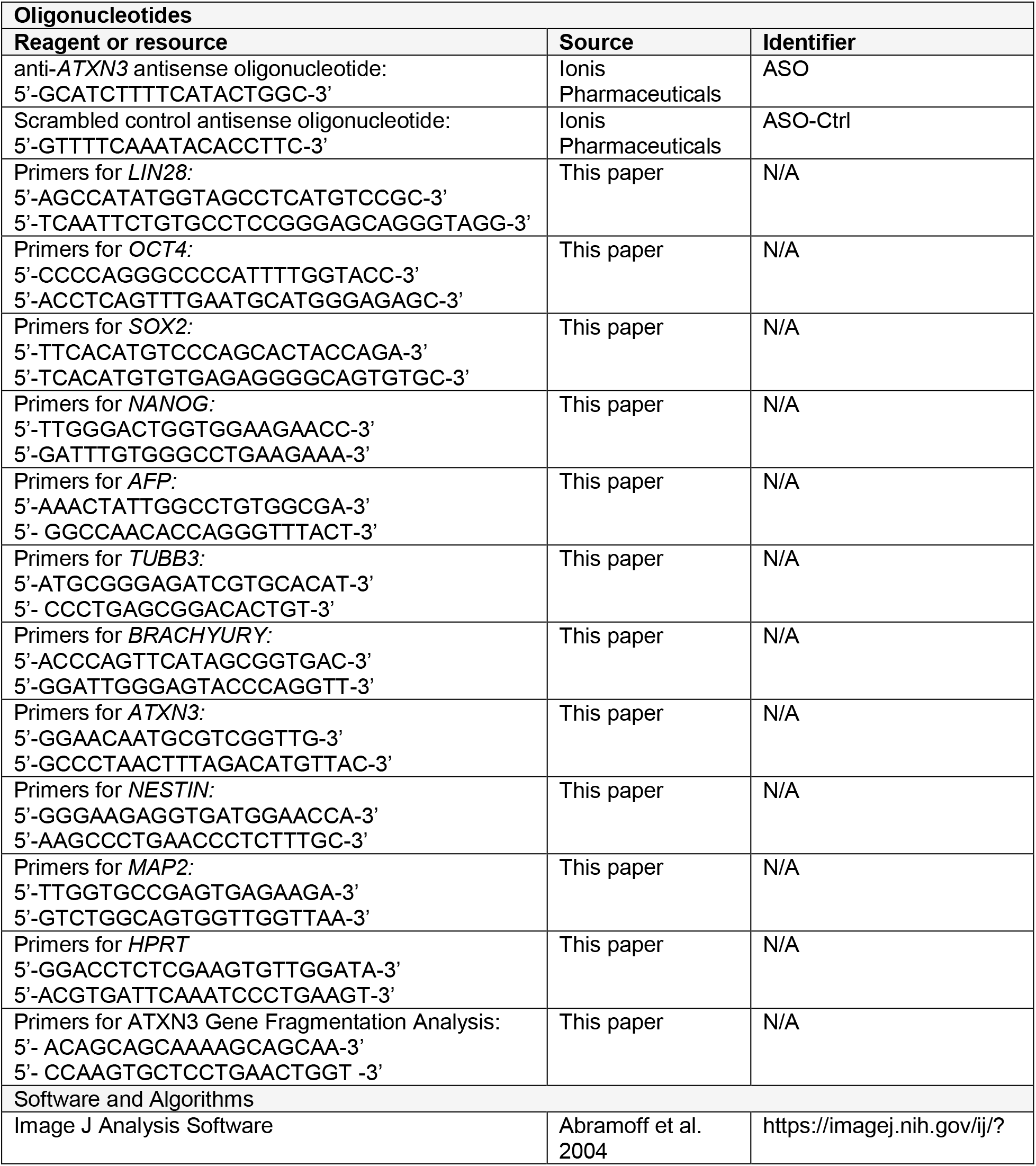

### 2.2 hESC Line Derivation and Characterization

Human embryos were originally created by assistive reproductive technologies for the purpose of procreation and donated to the University of Michigan under MStem Cell Laboratory’s Institutional Review Board (IRB) approved study *“Derivation of human Embryonic Stem Cells”* (HUM00028742). Written informed consent was obtained for all embryo donations. Donation and derivation of NIH-approved, unaffected hESC line UM4-6 (Registration #-0147, referred to here as WT-hESC) was reported previously^22^. The SCA3-affected embryo was donated to the University of Michigan following single gene preimplantation genetic testing identified the embryo as heterozygous for a pathogenic *CAG*-repeat expansion in *ATXN3.* Derived UM134-1 hESC line (referred to here as SCA3-hESC) was accepted to the NIH registry of approved hESC lines on September 29, 2014 (Registration # −0286).

Derivation of UM134-1 and derivatives were performed with non-federal funds prior to acceptance to the NIH registry. Prior to NIH approval, Cell Line DNA Fingerprinting and cytogenetic analysis were performed on passage 6 SCA3-hESC (Cell Line Genetics, Madison, Wisconsin). The resulting DNA STR profile of SCA3-hESC confirmed the presence of a single cell line that’s unique from lines published in the ATCC, NIH, or DSMZ websites. Cytogenetic analysis was performed on 20 G-banded metaphase cells. Seventeen cells demonstrated an apparently normal male 46,XY karyotype, while three cells demonstrated non-clonal aberrations ruled as most likely artifacts of culture. To determine *ATXN3* CAG-trinucleotide repeat length in WT- and SCA3-hESC, 10-50 μL of at least 10 ng/μL DNA were analyzed by gene fragmentation analysis (Laragen Inc., Culver City). Repeat length was calculated as (amplicon fragment size – 66) / 3.

### 2.3 Culture and ASO Transfection of hESC

Undifferentiated pluripotent hESC were cultured in mTeSR1 media (Stem Cell Technology) on Matrigel-coated plates with daily media changes and passaged using L7 passaging media (Lonza) or ReLeSR passaging media (Stem Cell Technology). The anti-ATXN3 ASO and scrambled control ASO (ASO-Ctrl) used for hESC transfections are 18 nucleotides in length with a mixed phosphodiester and phosphorothioate backbone and five MOE-modified nucleotides on each of the 5’ and 3’ termini. ASO nucleotide sequences are listed in the Key resources table. Oligonucleotides were synthesized as described previously^23,24^. ASOs were solubilized in PBS (without Ca^2+^ or Mg^2+^). For ASO transfections, undifferentiated hESC were plated on Matrigel-coated plates in mTeSR1 media supplemented with 10 μM Rock Inhibitor and grown overnight. Media was replaced with mTeSR1 the next day and cells were allowed to recover for at least 4 hours. Transfections were performed using TransIT LT1 transfection reagent (Mirus Bio) according to the manufacturer’s protocol. ASO concentration. Cells were cultured with daily media changes and harvested 3 days after transfection for RNA isolation or 4 days after transfection for Western blot analysis or immunocytochemical analysis.

### 2.4 Differentiation and Culture of NPCs and Forebrain CNs

Directed differentiation of hESC to NPCs and CNs was performed as described by Shi *et al.* (*Nature* 2012) with some modifications^25^. Briefly, undifferentiated hESC were cultured to confluency in mTeSR1 media (Stem Cell Technology) on Matrigel-coated plates. Neural differentiation was induced using neural maintenance media supplemented with 1 μM Dorsomorphin and 10 μM SB431542. Cells were cultured for approximately 12 days with daily media changes. Neuroepithelial cells were passaged onto Matrigel-coated plates using Dispase passaging reagent and were maintained in neural maintenance media containing 20 ng/ml FGF2 with daily media changes. Once neural rosettes were visible, NPCs were either collected for experimentation or cultured in neural expansion media supplemented with 20 ng/ml FGF and 20 ng/ml EGF and passaged as needed with Accutase passaging reagent. For forebrain CN differentiation, NPCs were plated onto PLO-Laminin coated plates or coverslips with neural expansion media for 24 hours, and then maintained in neural maintenance for 30 days with media changes every other day.

### 2.5 RNA Isolation and Quantitative PCR

Total RNA was isolated from cell lysates using Trizol reagent according to the manufacturer’s protocol (Invitrogen, Carlsbad, CA). Reverse transcription was performed on 0.5-1μg of total RNA using the iScript cDNA synthesis kit according to the manufacturer’s instructions (Bio-Rad, Hercules, CA). The cDNA was diluted 1:10 in nuclease-free water. iQ SYBR green quantitative polymerase chain reaction (qPCR) was performed on the diluted cDNA following the manufacturer’s protocol (Bio-Rad). Average-adjusted relative quantification analysis was performed using human primers listed in Key resources table.

### 2.6 Immunoblotting

Cell protein extracts were prepared from cells using two protocols. For immunoblot analysis of total protein extracts, cells were washed with cold PBS and lysed on ice with RIPA buffer containing complete mini protease inhibitor cocktail (Roche), followed by bath ultrasonication in chill water and centrifugation for total protein extract. For immunoblot analysis on soluble versus insoluble protein extracts, cells were washed and collected in PBS, bath ultrasonicated in chill water, centrifuged, and supernatants were collected (soluble fraction). Remaining pellets were washed with PBS and resuspended in 1% sarkosyl in PBS containing complete mini protease inhibitor cocktail (Roche), vortexed for 30 sec and incubated at room temperature for 1 hr. Samples were bath ultrasonicated again for 5 min, centrifuged, and supernatants were collected (insoluble fraction). Total protein concentrations were determined using the bicinchoninic acid (BCA) method (Pierce) and stored at −80°C. A total of 5 or 10 μg protein lysates were resolved in Novex 4-20% Tris-Glycine polyacrylamide electrophoresis gels (XP04205BOX; Invitrogen, Carlsbad, CA) and transferred to polyvinylidene difluoride (PVDF) membranes. Membranes were incubated overnight at 4°C with various antibodies listed in Key resources table. Bound primary antibodies were visualized by incubation with peroxidase-conjugated anti-mouse or anti-rabbit secondary antibody (1:10000; Jackson ImmunoResearch Laboratories, West Grove, PA) followed by treatment with the ECL-plus reagent (Western Lighting; PerkinElmer, Waltham, MA) and exposure to autoradiography films. Band intensities were quantified using ImageJ analysis software (NIH, Bethesda, MD). Band intensities for proteins of interest were divided by GAPDH or α-Tubulin band intensity, averaged, normalized to the mean control value indicated for each experiment, and expressed as % indicated control.

### 2.7 Immunocytochemistry and Image Analysis

Cells were cultured on Poly-D-Lysine/Matrigel-coated coverslips. Cells were washed with cold PBS and then fixed with 4% paraformaldehyde (PFA) solution for 15 min on ice. Cells were permeabilized with 0.1% Triton X-100 for 5 min at room temperature then blocked in a 5% Normal Goat Serum solution for 1 hr at room temperature. Cells were blocked in various primary antibodies overnight at 4°C. The cells were then immunostained with species-specific secondary antibodies conjugated to Alexa 488 or 568 fluorophores and mounted using Prolong Gold. Imaging was performed using an IX71 Olympus inverted microscope (Melville, NY) or Nikon-A1 confocal microscope (Melville, NY). All image analysis was performed using ImageJ analysis software (NIH, Bethesda, MD).

### 2.8 Electrophysiology

Electrophysiological recordings were performed on WT- and SCA3-CN plated on laminin-coated coverslips at 30-days post-differentiation. Cultured WT-or SCA3-CN were placed into a bath of room temperature Tyrode’s solution^26^ containing the following (in mmol/L): 82 Na_2_SO_4_, 30 K_2_SO_4_, 5 MgCl_2_, 10 HEPES, 10 glucose, at pH 7.4. Cells were slowly perfused with room-temperature Tyrode’s solution bubbled with 5% CO_2_ and 95% O_2_ (carbogen) during recordings. Patch-clamp recordings were performed with borosilicate glass pipettes of 4-5 mΩ resistance which contained an internal solution containing the following (in mmol/L): 119 K-gluconate, 2 Na-gluconate, 6 NaCl, 2 MgCl_2_, 0.9 EGTA, 10 HEPES, 14 Tris-phosphocreatine, 4 MgATP, 0.3 tris-GTP, at pH 7.3 and osmolarity 287 mOsm. An agarose bridge (1% in Tyrode’s solution) was used to shield cells from the AgCl bath electrode during recordings. Patch-clamp recordings were performed using an Axopatch 200B amplifier, Digidata 1440A interface, and pClamp-10 software (MDS Analytical Technologies, Sunnyvale, CA). Data were acquired at 100 kHz in the fast-current clamp mode of the amplifier and filtered at 2 kHz. Spontaneous neuronal activity was recorded in whole-cell current clamp mode to observe characteristic action potential shape. Whole-cell voltage clamp recordings were used in order measure currents. WT- and SCA3-CN were held at −80 mV, and peak inward and outward currents were quantified in response to depolarizing current steps from −90 mV to +10 mV in +10 mV increments. Series resistance was compensated up to 40%. Cells were included if series resistance remained under 30 MΩ for the duration of the recording.

### 2.9 Statistics

All statistical analyses were performed using Prism (7.0; Graph-Pad Software, La Jolla, CA). Quantified data represent the findings of three or more independent experiments. The statistical tests used are described in each figure legend. All error bars indicate variability about the mean expressed as standard error of the mean. Quantification of qRT-PCR transcript levels is normalized to mean WT-hESC, WT-NPC, or WT-CN transcript levels set at 1. Quantification of immunoblotted protein expression is normalized to mean WT-hESC, WT-NPC or WT-CN protein levels set at 100%. All tests set the level of significance at *p* < 0.05.

### 2.10 Key resources table

Key resource information is provided in a separate table.

## 3. RESULTS

### 3.1 Derivation and characterization of SCA3-hESC line UM134-1

The SCA3-affected embryo was originally created by assistive reproductive technologies for the purpose of procreation. The embryo was donated to the University of Michigan following single gene preimplantation genetic testing, which identified the embryo as heterozygous for a pathogenic *CAG* repeat expansion in *ATXN3*. The derived UM134-1 hESC line (referred to here as SCA3-hESC) was accepted to the NIH registry of approved hESC lines in 2014 (#0286). The unrelated hESC line UM4-6 (#0147, referred to here as WT-hESC), developed previously under the same protocol at the University of Michigan and also accepted to the NIH registry, was used throughout these studies as a comparative control^22^.

Undifferentiated SCA3-hESC exhibited a cellular morphology characteristic of pluripotent stem cells and express pluripotency markers OCT3/4, SOX2 and NANOG (Figure 1A). Quantitative real-time PCR (qRT-PCR) confirmed that SCA3-hESC express pluripotency markers *LIN28*, *OCT4*, *SOX2*, and *NANOG* transcripts at levels comparable to WT-hESC (Figure 1B) (n = 3 passages). Following 21 days of differentiation, PCR analysis confirmed that SCA3-hESC derived embryoid bodies express lineage markers *AFP* (endoderm), *BRACHYURY* (mesoderm), and *TUJ-1* (ectoderm) (Figure 1C). G-banded cytogenetic analysis of SCA3-hESC identified a normal 46,XY karyotype with no chromosomal abnormalities (Figure 1D). A previous study showed that the WT-hESC line used here also has a normal 46,XY karyotype with no chromosomal abnormalities^22^.

**Figure 1.**
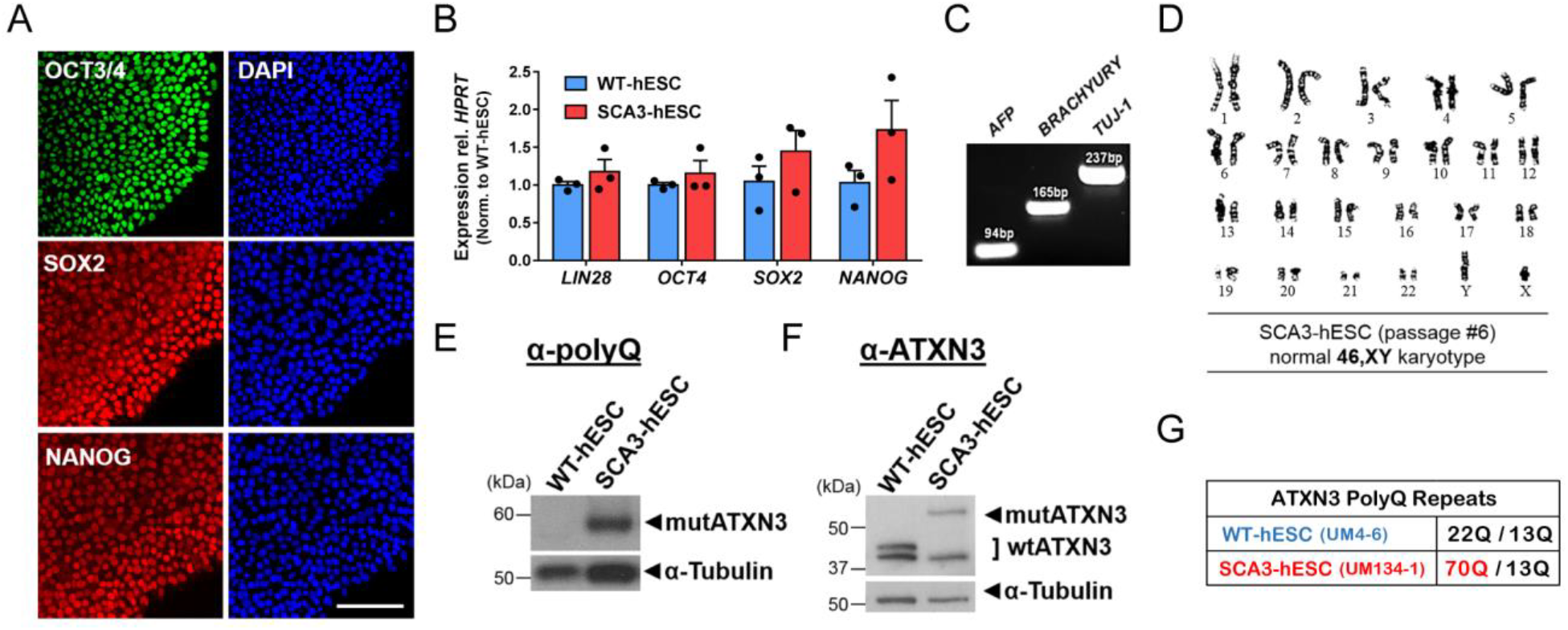
SCA3-hESC line UM134-1 is pluripotent, possesses a normal karyotype, and expresses pathogenic polyglutamine-expanded mutant ATXN3. (A) Immunocytochemical analysis with DAPI co-stain of SCA3-hESC revealed expression of pluripotency markers OCT3/4, SOX2, and NANOG. Scale bar = 200 μm. (B) Undifferentiated WT-hESC (blue) and SCA3-hESC (red) expressed pluripotency markers *LIN28*, *OCT4*, *SOX2* and *NANOG* as assessed by qRT-PCR analysis. Data are represented as mean of three replicates ± SEM. (C) SCA3-hESC were differentiated into embryoid bodies for 21 days in culture. Differentiated SCA3-embryoid bodies expressed lineage markers of endodermal [*α-fetoprotein* (*AFP*)], mesodermal [*Brachyury*], and ectodermal tissue [*neuron-specific class III beta-tubulin* (*TUJ-1*)]. Electrophoresis demonstrated anticipated amplicon size for each lineage marker PCR primer set. (D) G-banded karyotype analysis of passage 6 undifferentiated SCA3-hESC showed a normal 46,XY karyotype. (E) Representative anti-polyQ expansion and (F) anti-ATXN3 Western blot of undifferentiated WT- and SCA3-hESC revealed heterozygous expression of polyQ-expanded ATXN3 protein in SCA3-hESC within the pathogenic range for SCA3. (G) Anticipated ATXN3 polyQ repeat lengths in WT- and SCA3-hESC lines as determined by gene fragmentation analysis. Mutant polyQ-expanded repeat length is highlighted in red. (mutATXN3 = mutant ATXN3; wtATXN3 = wild type ATXN3).

Immunoblot analysis of WT- and SCA3-hESC protein extracts using monoclonal antibody 1C2, which recognizes expanded polyQ, revealed a band of ~60 kDa in SCA3-hESC that is not present in WT-hESC, indicating expression of a polyQ-expanded protein in SCA3-hESC (Figure 1E). Anti-ATXN3 immunoblot analysis identified a doublet of ~42-44 kDa in WT-hESC consistent with normal repeat length polymorphism in *ATXN3*, whereas ATXN3 bands of ~42kDa and ~60kDa were observed in SCA3-hESC, corresponding to wild type ATXN3 (wtATXN3) and polyQ-expanded mutant ATXN3 (mutATXN3), respectively (Figure 1F). Subsequent gene fragmentation analysis performed on early passages of both lines confirmed that SCA3-hESC is heterozygous for one pathogenic and one normal *ATXN3* allele encoding ATXN3 polyQ expansions of 70Q and 13Q, respectively, compared to non-pathogenic WT-hESC polyQ lengths of 22Q and 13Q (Figure 1G).

### 3.2 Undifferentiated SCA3-hESC exhibit ATXN3 aggregation and nuclear sequestration

Misfolding of expanded polyQ proteins and subsequent accumulation as high molecular weight oligomers and aggregates is a key pathological feature of polyQ diseases^4,27,28^. SCA3 post-mortem patient brains and transgenic mouse models of SCA3 exhibit widespread neuronal intranuclear and cytosolic inclusions of ATXN3 in brain regions affected in disease^28–32^. Aggregation of polyQ-expanded mutATXN3 has been detected in cellular models, but to our knowledge only with the aid of genetic or environmental manipulations, such as overexpression of highly aggregate-prone ATXN3 fragments^33–35^ or stress induction through proteasomal inhibition or excitotoxicity^12,36^. A human cell model showing aggregation of mutATXN3 expressed at endogenous levels in the absence of exogenous stressors would be an excellent tool for advancing our understanding of the role of native ATXN3 misfolding and aggregation in SCA3 pathogenesis.

We first sought to determine whether SCA3-hESC recapitulated mutATXN3 misfolding and aggregation under normal culture conditions. Anti-ATXN3 immunoblot analysis of undifferentiated WT- and SCA3-hESC protein lysates demonstrated accumulation of high molecular weight (HMW) ATXN3 species in SCA3-hESC that was largely absent in WT-hESC (Figure 2A). Quantification confirmed an 80% ± 21% increase in HMW ATXN3 species >100kDa in SCA3-hESC relative to WT-hESC (*p* = 0.02, n = 3 passages) (Figure 2B). Anti-ATXN3 immunocytochemistry (ICC) with DAPI co-staining revealed striking intranuclear and cytoplasmic ATXN3-positive puncta of variable sizes and shapes in SCA3-hESC (Figure 2C). Though rare puncta were observed in WT-hESC, SCA3-hESC formed significantly more puncta, averaging ~1 per cell with many cells containing multiple puncta (SCA3-hESC = 0.93 ± 0.12 ATXN3 puncta/cell, WT-hESC = 0.04 ± 0.01 ATXN3 puncta/cell, *p*<0.0001, n = 3-4 passages, 3-5 confocal images/passage) (Figure 2C and 2D).

**Figure 2.**
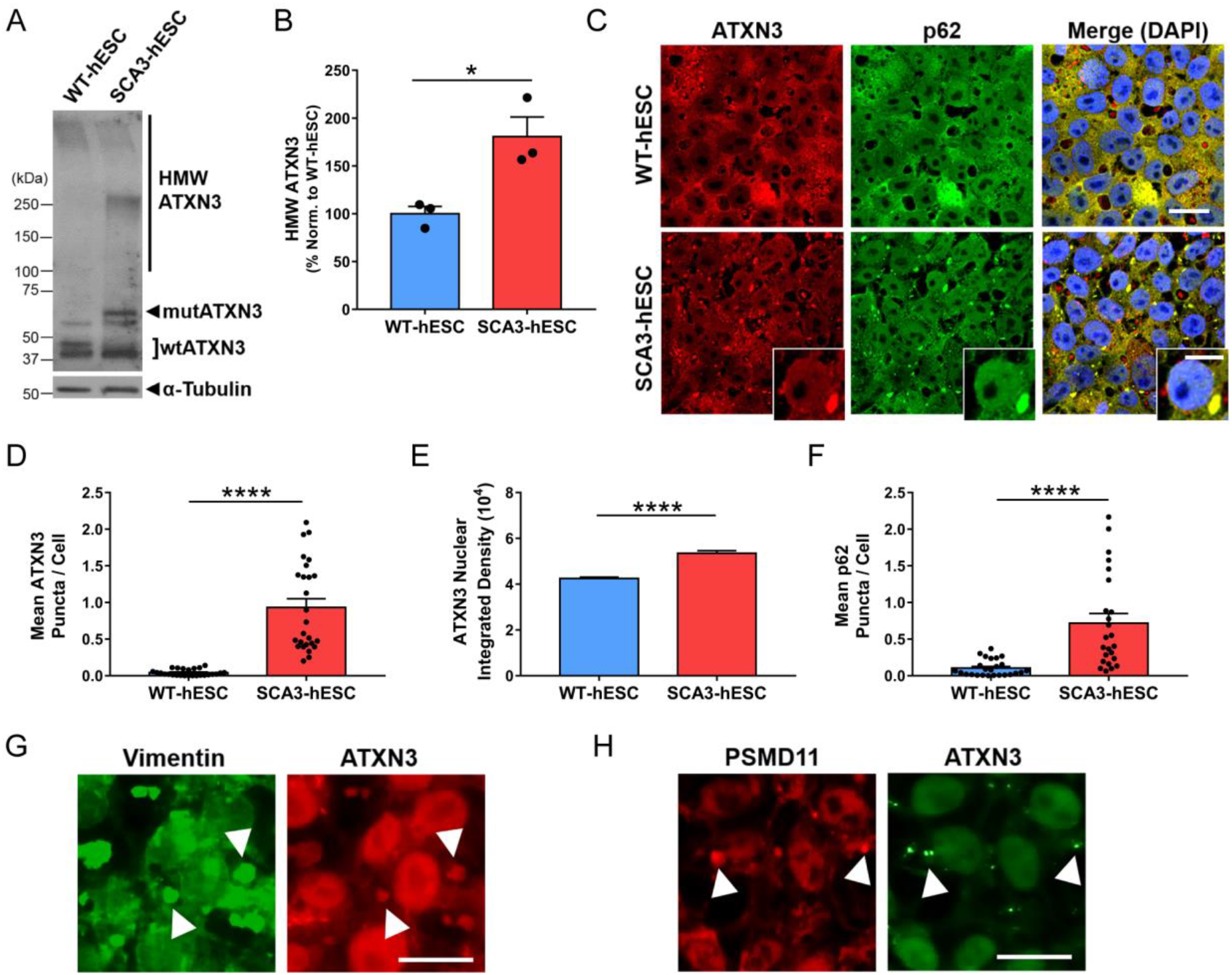
SCA3-hESC form high molecular weight ATXN3 aggregates that localize to p62-positive aggresomes, and exhibit enhanced nuclear sequestration of ATXN3. (A) Representative anti-ATXN3 Western blot and (B) quantification of high molecular weight (HMW) ATXN3 species in undifferentiated WT-hESC (blue) and SCA3-hESC (red). Data (mean of three replicates ± SEM) are reported normalized to WT-hESC set to 100%. (C) Immunocytochemistry of undifferentiated WT- and SCA3-hESC showing single channel anti-ATXN3 (red) and anti-p62 (green) immunofluorescence merged with DAPI co-stain (blue). Scale bar = 25 μm. Inset scale bar = 10 μm. (D) Mean ATXN3 puncta per cell, (E) ATXN3 nuclear integrated density, and (F) p62 puncta per cell in WT- and SCA3-hESC. (G) Anti-vimentin (green) and anti-ATXN3 (red) immunofluorescence in undifferentiated SCA3-hESC. Scale bar = 20 μm. (H) Anti-PSMD11 (red) and anti-ATXN3 (green) immunofluorescence in undifferentiated SCA3-hESC. White arrowheads indicate co-localization of vimentin^+^ or PSMD11^+^ and ATXN3^+^ puncta. Scale bar = 20 μm. Data (mean ± SEM) are reported (n = 3-5 confocal images per 3-4 independent replicates). Data points in (D) and (F) represent individual confocal images, while mean nuclear integrated density in (E) was calculated by averaging across imaged nuclei. Unpaired two-tailed t test (\**p*<0.05, \*\**p*<0.01, \*\*\**p*<0.001, \*\*\*\**p*<0.0001). (mutATXN3 = mutant ATXN3; wtATXN3 = wild type ATXN3; PSMD11 = proteasome 26S subunit, non-ATPase 11).

Abnormal sequestration of ATXN3 into neuronal nuclei is also a prominent pathological feature of SCA3 human disease^28,37–39^. Possessing both nuclear export and nuclear localization sequences, non-pathogenic ATXN3 is normally diffusely localized throughout the cytosol and nucleus^39^, but undergoes rapid nuclear translocation following induction of proteotoxic stress^36^. Though it remains unclear why polyQ-expanded mutATXN3 becomes sequestered in neuronal nuclei, preventing or decreasing nuclear import of mutATXN3 rescues disease in SCA3 transgenic mouse models, suggesting that nuclear localization of mutATXN3 is a necessary step in disease pathogenesis^37^. To determine whether SCA3-hESC replicate this key pathological hallmark of disease, we quantified mean ATXN3 nuclear integrated density in undifferentiated WT- and SCA3-hESC, employing anti-ATXN3 and DAPI co-stained confocal imaging. Subsequent ImageJ analysis revealed an over 25% increase in mean ATXN3 nuclear integrated density in SCA3-hESC relative to WT-hESC (*p* < 0.0001) (Figure 2C and 2E). Thus, in contrast to previously reported SCA3 *in vitro* models^12,33–36^, SCA3-hESC endogenously form nuclear and cytoplasmic ATXN3 inclusions and exhibit enhanced nuclear sequestration of ATXN3 under normal, unstressed culture conditions.

### 3.3 Large ATXN3 aggregates localize to p62^+^ aggresomes in SCA3-hESC

To further characterize observed ATXN3^+^ puncta, we investigated whether proteins typically co-localizing with aggregates in SCA3 patient brains similarly co-localize in SCA3-hESC. The ubiquitin-binding shuttle protein p62 is found in ubiquitinated protein aggregates in many neurodegenerative diseases^40^, including ATXN3 aggregates in SCA3 patient post-mortem brains^30,41,42^. Confocal imaging was performed on undifferentiated WT- and SCA3-hESC following anti-p62 and anti-ATXN3 immunofluorescence and DAPI staining (Figure 2C). Unlike WT-hESC, in which p62 was more diffuse throughout cells and formed few p62+ puncta, SCA3-hESC formed large, juxtanuclear p62^+^ puncta: SCA3-hESC = 0.72 ± 0.13 p62 puncta/cell; WT-hESC = 0.10 ± 0.02 p62 puncta/cell (*p* < 0.0001) (Figure 2C and 2F). Large p62^+^ structures always co-localized with large ATXN3^+^ puncta (Figure 2C), although many ATXN3^+^ puncta lacked p62-staining suggesting there are two or more ATXN3 inclusion states in SCA3-hESC. In addition, p62 exhibited significantly increased localization to the nucleus in SCA3-hESC (mean p62 nuclear integrated density: SCA3-hESC = 58065 ± 1141; WT-hESC = 45332 ± 743, *p* <0.0001). Previous studies have shown that inhibiting autophagy induces p62 nuclear sequestration, thus enhanced nuclear p62 may indicate autophagic impairment in SCA3-hESC^43^.

Notably, p62 promotes the formation of mutATXN3 aggresomes in cell-based overexpression model systems^44^. Aggresomes are dynamic pericentriolar structures consisting of a condensed protein aggregate core contained within an intermediate filament vimentin cage that function as a cellular pathway to consolidate misfolded protein aggregates^45^. Anti-vimentin and anti-ATXN3 ICC was performed on WT- and SCA3-hESC to investigate the possibility of aggresome formation in SCA3-hESC. SCA3-hESC formed distinct vimentin^+^ puncta that consistently coincided with large ATXN3^+^ puncta (Figure 2G). Like p62, vimentin^+^ puncta were rarely observed in WT-hESC, and when present did not preferentially co-localize with ATXN3 (data not shown). Further ICC investigations revealed that the 26S proteasome subunit PSMD11 also selectively localized to ATXN3 puncta in SCA3-hESC, demonstrating that components of the ubiquitin-dependent protein quality control machinery are also being sequestered into aggregates in SCA3-hESC (Figure 2H). Together, these results suggest SCA3-hESC, but not WT-hESC, form ATXN3 aggregates, the majority of which are consolidated into p62^+^ aggresomes.

### 3.4 Altered expression of key protein clearance pathway proteins in SCA3-hESC

Aggresome formation suggests a failure of canonical degradation pathways to dispose of misfolded proteins^45^. To determine which pathways may contribute to aggresome formation in SCA3-hESC, we evaluated key proteins involved in protein homeostasis including markers for autophagy, ubiquitin-dependent proteasomal degradation and protein folding. The autophagy-linked protein p62 is selectively degraded through autophagic clearance and thus increased levels of p62 may indicate autophagic inhibition^46^. Immunoblot analysis showed that p62 levels were increased by 80% ± 24% in SCA3-hESC compared to WT-hESC (*p* = 0.008, n = 3 passages per line, 2-3 replicates per passage) (Figure 3A and 3B). HMW p62, which is presumably aggregated, also accumulated in SCA3-hESC protein lysates, likely corresponding to p62 interactions with insoluble aggresome complexes formed by mutATXN3 (Figure 3A and 3B). The expression of autophagy-related proteins Beclin1 and LC3-II was also compared in WT- and SCA3-hESC protein lysates (Figure 3C). Though variable, LC3-II protein levels were significantly decreased by 64% ± 26% in SCA3-hESC relative to WT-hESC (*p* = 0.037), whereas Beclin1 levels did not statistically differ (*p* = 0.57) (n = 3 passages per line, 2-3 replicates per passage) (Figure 3D).

**Figure 3.**
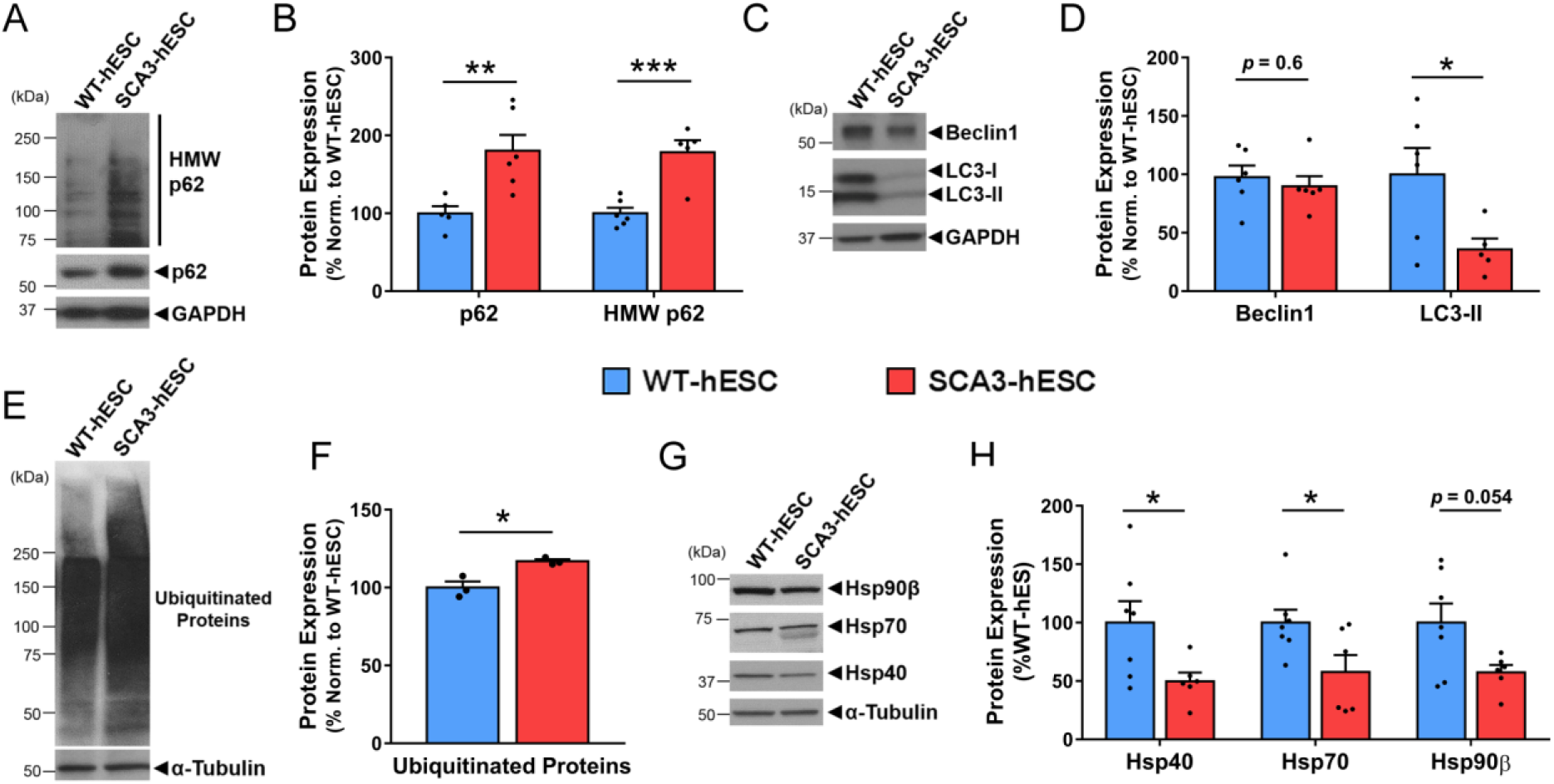
SCA3-hESC exhibit altered expression of key regulators of protein homeostasis. (A) Representative anti-p62 Western blot and (B) quantification of monomeric and high molecular weight (HMW) p62 in undifferentiated WT-hESC (blue) and SCA3-hESC (red) (n = 6). (C) Representative anti-Beclin1 and anti-LC3B Western blot and (D) quantification in undifferentiated WT- and SCA3-hESC (n = 6). (E) Representative anti-ubiquitin western blot and (F) quantification of ubiquitinated proteins in WT- and SCA3-hESC (n = 3). (G) Anti-Hsp40, anti-Hsp70, and anti-Hsp90β Western blots and (H) quantification in WT- and SCA3-hESC (n = 6). Data (mean of 3-6 independent replicates ± SEM) are reported relative to WT-hESC set to 100%. Unpaired two-tailed t test (\**p*<0.05, \*\**p*<0.01, \*\*\**p*<0.001). (GAPDH = glyceraldehyde-3-phosphate dehydrogenase).

Increased levels of ubiquitinated substrates have been observed in human SCA3 brain tissue and mouse models of SCA3, possibly reflecting altered mutATXN3 deubiquitinase activity^47^ and downstream impairment in proteolytic clearance pathways^48,49^. Using a pan-ubiquitin antibody, we found that poly-ubiquitinated protein levels were slightly but significantly increased by 17% ± 4% in SCA3-hESC relative to WT-hESC (*p* = 0.015, n = 3 passages per line) (Figure 3E and 3F), suggesting impairment in ubiquitin-dependent proteasomal clearance. Heat shock proteins Hsp40, Hsp70, and Hsp90β, which play important roles in maintaining protein homeostasis in cells^50^, are also altered in SCA3 post-mortem brains and disease model systems^50–53^. Immunoblot analysis revealed Hsp40 and Hsp70 to be significantly decreased by 50% ± 21% (*p* = 0.037) and 42% ± 18% (*p* = 0.039), respectively, in SCA3-hESC relative to WT-hESC (n = 3 passages per line, 2-3 replicates per passage) (Figure 3G and 3H). SCA3-hESC also consistently exhibited a doublet band for Hsp70 that was absent from WT-hESC, which could indicate altered post-translational modifications, splicing, or protein-protein interactions of Hsp70 in SCA3-hESC. Hsp90β also trended towards decreased levels in SCA3-hESC, but failed to reach statistical significance (*p* = 0.054) (Figure 3G and 3H). Though further analysis is required to determine the extent and exact nature of autophagic and proteasomal dysfunction in SCA3-hESC, the altered levels of key regulators of protein quality control spanning autophagy, ubiquitin-dependent proteasomal degradation and protein folding machinery suggest widespread disruption of protein clearance in SCA3-hESC, a characteristic feature of SCA3 pathogenesis^1^.

### 3.5 ASO-mediated ATXN3 reduction rescues aggresome formation in SCA3-hESC

To assess the potential of SCA3-hESC as a cellular model for preclinical therapeutic testing, we investigated whether delivery of an anti-*ATXN3* antisense oligonucleotide (ASO) recently validated in preclinical studies^20,21^ could rescue molecular phenotypes in SCA3-hESC. ASOs are short, chemically modified nucleotide sequences that, when bound to sequence-specific target mRNA, activate RNaseH-mediated degradation of the targeted mRNA thereby reducing expression of the encoded protein^54^. In preclinical studies, we recently demonstrated that a single intracerebroventricular injection of an anti-*ATXN3* ASO into a transgenic mouse model of SCA3 suppressed expression of mutATXN3, reduced neuronal nuclear ATXN3 accumulation, and rescued several molecular, electrophysiological, and behavioral phenotypes in SCA3 mice^20,21^.

WT- and SCA3-hESC were transfected with 100 nM or 500 nM anti-*ATXN3* ASO (referred to previously as ASO-5^20,21^), a non-specific ASO control (ASO-Ctrl) or PBS vehicle, then harvested three or four days later for RNA and protein analysis, respectively. Immunoblot analysis identified a significant decrease in both wtATXN3 and mutATXN3 protein levels in SCA3-hESC transfected with 100 nM and 500 nM compared to vehicle-treated or ASO-Ctrl treated SCA3-hESC groups [500 nM: wtATXN3 = 63% ± 8%, (*p* = 0.0003); mutATXN3 = 70% ± 7% (*p* = 0.0015)], normalized to vehicle-treated SCA3-hESC set at 100%, n = 6 replicates per treatment group] (Figure 4A). qRT-PCR analysis confirmed an approximately 75% reduction in total *ATXN3* transcript levels in SCA3-hESC transfected with 500 nM ASO relative to vehicle-treated SCA3-hESC (*p* = 0.03, n = 3) (Figure 4B). SCA3-hESC transfected with 100 nM ASO trended towards decreased *ATXN3* expression levels at 72 hours, but failed to reach significance (*p* = 0.09, n = 3 replicates per treatment group) (Figure 4B).

**Figure 4.**
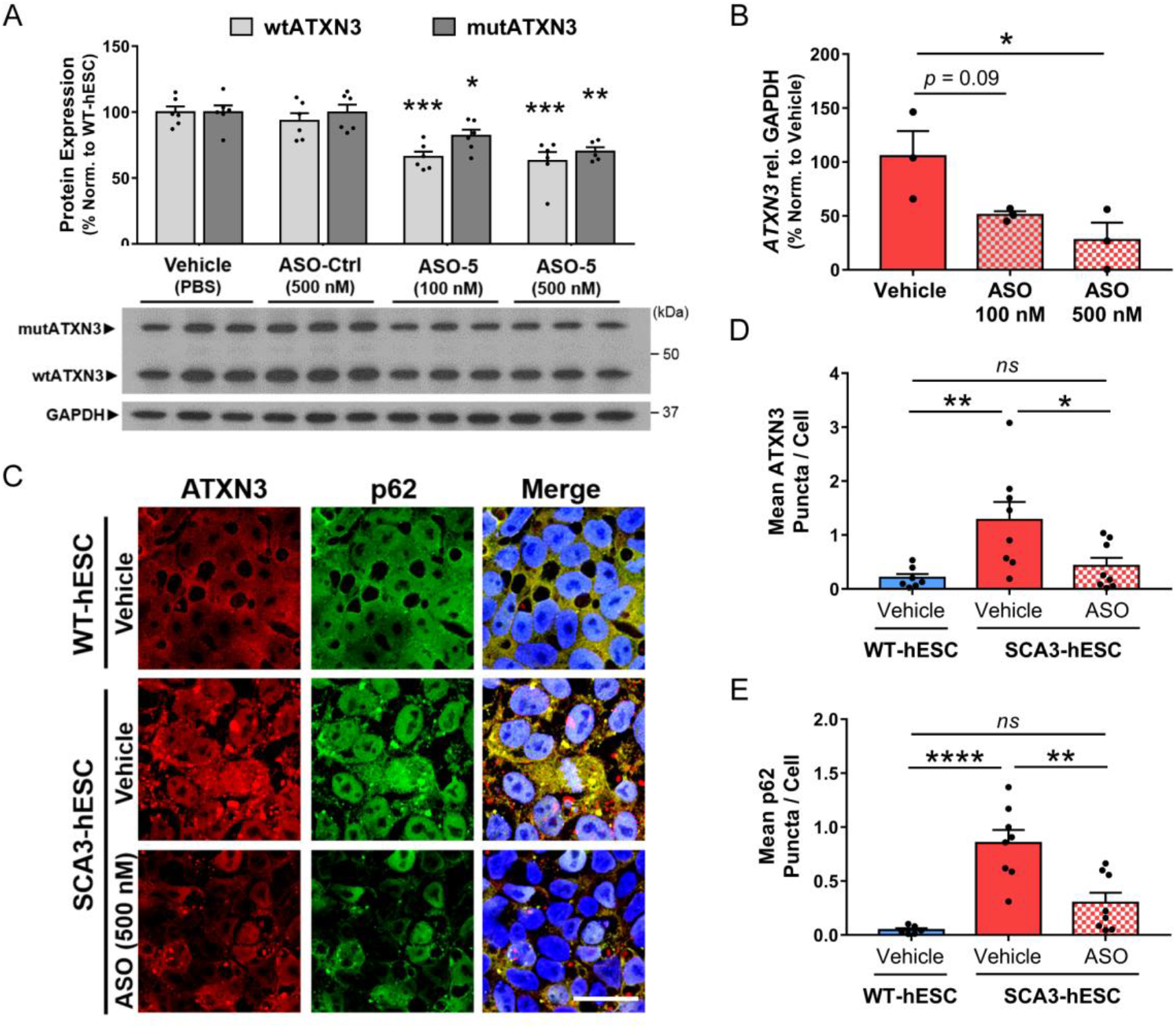
Anti-*ATXN3* antisense oligonucleotide-mediated reduction of ATXN3 rescues ATXN3 aggregation and aggresome formation in SCA3-hESC. (A) Representative anti-ATXN3 Western blot and quantification of mutant ATXN3 (mutATXN3, dark gray) and wild type ATXN3 (wtATXN3, light gray) in SCA3-hESC protein lysates four days after transfection with PBS vehicle, 500 nM of scrambled antisense oligonucleotide control (ASO-Ctrl), and 100 nM or 500 nM of an anti-*ATXN3* antisense oligonucleotide (ASO) (n = 6 replicates). (B) qRT-PCR analysis of total *ATXN3* transcript levels in SCA3-hESC three days after transfection vehicle, 100 nM or 500 nM of ASO (n = 3 replicates). (C) Anti-ATXN3 (red) and anti-p62 (green) immunofluorescence and DAPI co-stain (blue) merged images in vehicle-treated WT-hESC, vehicle-treated SCA3-hESC, and SCA3-hESC transfected with 500 nM ASO. Cells were fixed four days following transfection of vehicle or ASO. Scale bar = 25 μm. (D) Quantification of mean ATXN3 puncta per cell and (E) mean p62 puncta per cell in WT-hESC (blue) and SCA3-hESC (red) transfected with vehicle, and SCA3-hESC transfected with 500 nM anti-ATXN3 ASO (red/white checkered) (n = 2-3 images quantified from 3 experimental replicates). Data (mean ± SEM) are reported. One-way ANOVA performed with the post-hoc Tukey test. (\**p*<0.05, \*\**p*<0.01, \*\*\*\**p*<0.0001). (GAPDH = glyceraldehyde-3-phosphate dehydrogenase; *ns =* not significant).

To determine whether ASO treatment rescued ATXN3 aggregation and p62^+^ aggresome formation, we performed anti-ATXN3 and anti-p62 ICC on SCA3-hESC transfected with 500 nM ASO compared to vehicle-treated WT- and SCA3-hESC (Figure 4C). Four days after transfection, diffuse and punctate ATXN3 fluorescence was visibly reduced in ASO-treated SCA3-hESC relative to vehicle-treated SCA3-hESC (Figure 4C). Importantly, ASO-mediated reduction in ATXN3 markedly reduced ATXN3 puncta per cell in SCA3-hESC to levels non-statistically different from that of WT-hESC (vehicle-treated WT-hESC = 0.21 ± 0.07 ATXN3 puncta/cell, vehicle-treated SCA3-hESC = 1.28 ± 0.33 ATXN3 puncta/cell, ASO-treated SCA3-hESC = 0.43 ± 0.15 ATXN3 puncta/cell, n = 3 replicates per treatment group, 2-3 confocal images per replicate) (Figure 4C and 4D). Similarly, ASO treatment led to visibly reduced diffuse p62 and completely rescued mean p62 puncta per cell four days after ASO transfection (vehicle-treated WT-hESC = 0.05 ± 0.02 p62 puncta/cell, vehicle-treated SCA3-hESC = 0.85 ± 0.12 p62 puncta/cell, ASO-treated SCA3-hESC = 0.30 ± 0.09 p62 puncta/cell, *p*<0.0001, n = 3 replicates per treatment group, 2-3 confocal images per replicate) (Figure 4C and 4E).

In summary, non-allele specific suppression of ATXN3 with an anti-*ATXN3* ASO rescued ATXN3 aggregation and p62^+^ aggresome formation in SCA3-hESC despite achieving a relatively modest 30% reduction in soluble mutATXN3 protein. These findings support the view that these disease-relevant molecular phenotypes are caused by expression of the pathogenic disease protein and do not reflect non-specific hESC line variations between WT- and SCA3-hESC. The identification of aggresome formation as a disease-dependent, reversible molecular phenotype bolsters its potential as a disease biomarker and the potential utility of this line for therapeutic preclinical testing for SCA3.

### 3.6 SCA3-hESC can be differentiated into neural progenitor cells and MAP2^+^ cortical neurons

As a degenerative brain disease^1^, SCA3 is thought principally to involve neurons. Thus, studying the disease in human neuronal cultures harboring the endogenous causative mutation should provide biologically relevant insights into disease. WT- and SCA3-hESC were differentiated into neural progenitor cells (NPC) and cortical neurons (CN) at similar efficiencies using a previously described protocol^25^. qRT-PCR analysis confirmed loss of pluripotency marker *OCT4* and gain of neural lineage marker *NESTIN* in derived NPC and 30-days post-differentiation (Day 30) CN, with similar *NESTIN* transcript levels achieved in both WT- and SCA3-NPC (Figure 5A and 5B). In addition, differentiated WT- and SCA3-CN expressed mature neuronal marker *MAP2* at similar levels 30-days post-differentiation (Figure 5C). ICC analysis confirmed expression of the neural lineage marker NESTIN in derived SCA3-NPC (Figure 5D) and the mature neuronal marker MAP2 in differentiated SCA3-CN (Figure 5E).

**Figure 5.**
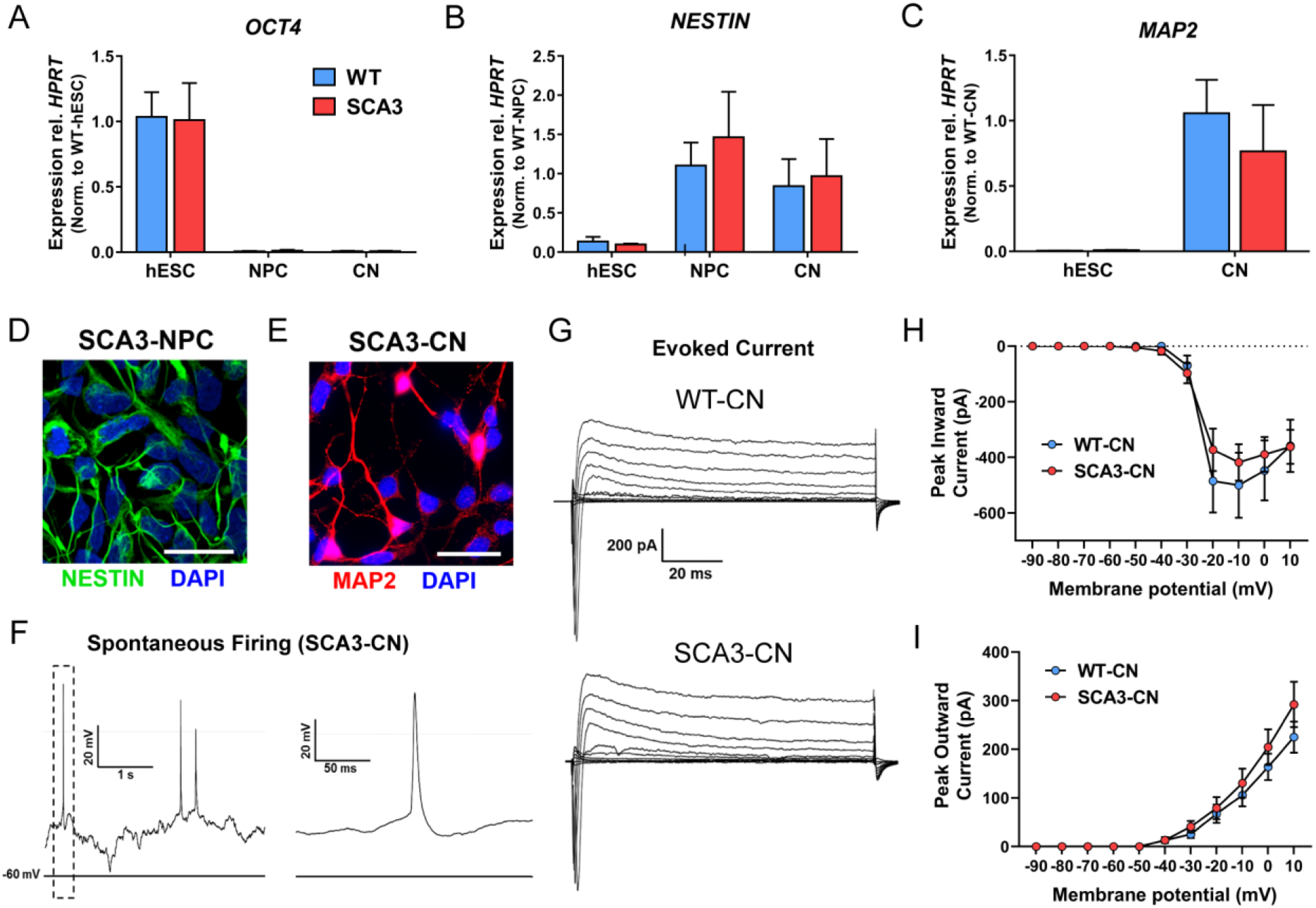
SCA3-hES can be differentiated into NESTIN^+^ neural progenitor cells and Day 30 MAP2^+^ forebrain cortical neurons that exhibit spontaneous firing activity. (A) qRT-PCR analysis for pluripotency marker *OCT4*, (B) neural stem cell marker *NESTIN*, and (C) mature neural lineage marker *MAP2* in undifferentiated WT-(blue) and SCA3-hESC (red) and differentiated neural progenitor cells (NPC) and cortical neurons (CN). Data (mean ± SEM) are reported normalized to WT-hESC, WT-NPC, or WT-CN as indicated in graph (n = 3 replicates). (D) NPCs derived from SCA3-hESC immunostained for neural stem cell marker NESTIN (green) and DAPI (blue). Scale bar = 50 μm. (E) Neurons derived from SCA3-hESC immunostained with mature neuronal marker MAP2 (red) and DAPI (blue). Scale bar = 25 μm. (F) Representative whole-cell current-clamp electrophysiological recording and enlarged inset of SCA3-CN showing spontaneous action potentials. (G) Representative whole-cell voltage-clamp recordings of WT- and SCA3-CN indicating evoked inward and outward currents in response to depolarizing voltage steps. (H) Quantification of peak inward currents in response to depolarizing voltage steps in WT-(n = 7 cells) and SCA3-CN (n = 9 cells). (I) Quantification of peak outward currents in response to depolarizing voltage steps in WT-(n=7 cells) and SCA3-CN (n = 9 cells). Data (mean ± SEM) are reported. Two-way ANOVA performed with Holm Sidak correction for multiple comparisons.

To further characterize neuronal maturation, we performed whole-cell electrophysiological recordings of Day 30 WT- and SCA3-CN. In current clamp mode, we observed spontaneous neuronal firing with characteristic action potential shape in SCA3-CN 30-days post-differentiation (Figure 5F). Voltage-clamp recordings of WT- and SCA3-CN indicated both inward and outward currents in response to depolarizing voltage steps with no statistical difference between WT- and SCA3-CN peak inward current [F(1,14)=0.19, *p*=0.67] or peak outward current [F(1,14)=0.66, *p*=0.43] (Figure 5G-5I). These data suggest proper maturation of derived WT- and SCA3-CN into MAP2-positive neurons that exhibit firing properties, with no indication of impaired neuronal differentiation of SCA3-hESC.

### 3.7 Robust ATXN3 aggregation in SCA3-hESC derived neural progenitor cells and cortical neurons

After confirming NPC and CN differentiation, we evaluated ATXN3 expression and aggregation propensity in derived SCA3-NPC and SCA3-CN. Soluble wtATXN3 and mutATXN3 were expressed in differentiated SCA3-NPC and Day 30 SCA3-CN at the expected molecular weights (Figure 6A and 6C). Similar to SCA3-hESC, SCA3-NPC and SCA3-CN exhibited markedly increased accumulation of HMW ATXN3. Quantified HMW ATXN3 levels were about 2.5 times greater in SCA3-NPC relative to WT-NPC (*p* < 0.001) (Figure 6A and 6B) and 1.7 times greater in SCA3-CNs relative to WT-CNs (*p* = 0.003) (Figure 6C and 6D).

**Figure 6.**
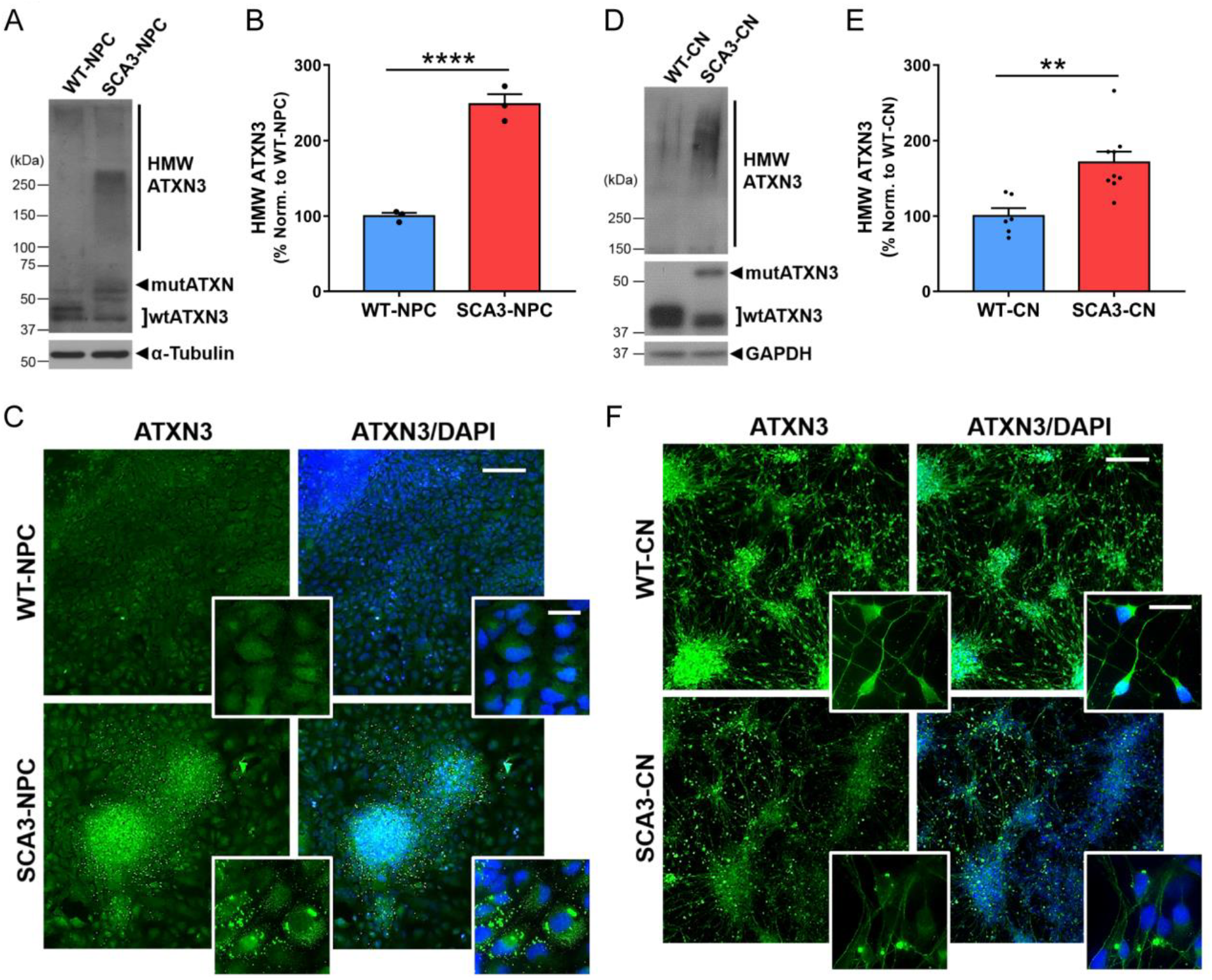
SCA3-hESC derived neural progenitor cells and Day 30 cortical neurons express polyglutamine-expanded mutant ATXN3 and accumulate high molecular weight ATXN3 aggregates. (A) Representative anti-ATXN3 Western blot and (B) quantification of high molecular weight (HMW) ATXN3 levels in neural progenitor cells (NPC) derived from WT-hESC (WT-NPC, blue) and SCA3-hESC (SCA3-NPC, red). Data (mean ± SEM) are reported relative to mean WT-NPC set to 100% (n=3 replicates). (C) WT- and SCA3-NPC immunostained for ATXN3 (green) and DAPI (blue). (D) Representative anti-ATXN3 Western blot and (E) quantification of HMW ATXN3 levels in cortical neurons (CN) derived from WT-hESC (WT-CN, blue) and SCA3-hESC (SCA3-CN, red). Data (mean ± SEM) are reported relative to mean WT-CN set to 100% (n=3 replicates). (F) WT-CN and SCA3-CN shown with anti-ATXN3 (green) and DAPI (blue) immunostaining. Scale bar = 200 μm. Inset scale bar = 25 μm. Unpaired two-tailed t test (\*\**p*<0.01, \*\*\*\**p*<0.0001).

We then performed anti-ATXN3 ICC with DAPI counter-stain on NPC and Day 30 CN differentiated from WT- and SCA3-hESC. Whereas ATXN3 was diffusely expressed throughout the nucleus and cytoplasm in WT-NPC, ATXN3 in SCA3-NPC localized to numerous cytoplasmic aggregates that varied in size, shape, number and location (Figure 6E). Differentiated SCA3-CN also exhibited robust ATXN3 inclusion formation, with individual neurons frequently forming one or more large, round, juxtanuclear ATXN3-positive inclusions as well as occasional round inclusions in distal processes (Figure 6F). By contrast, inclusions were rarely observed in WT-NPC or WT-CN. Unlike the case in SCA3-hESC, enhanced nuclear accumulation of ATXN3 was not observed in SCA3-NPC or SCA3-CN. These data demonstrate robust aggregation when mutATXN3 is expressed in SCA3-hESC, -NPC, and -CN under normal culture conditions in the absence of applied stressors, which has not been described in any other SCA3 disease-specific iPSC or hESC line.

### 3.8 Altered expression of key protein clearance pathway proteins in SCA3-CN

We also analyzed whether the dysregulation of components of the protein homeostasis machinery observed in undifferentiated SCA3-hESC persisted in Day 30 SCA3-CN. WT- and SCA3-CN were fractionated into Triton-X100 soluble and insoluble protein extracts which were then subjected to immunoblot analysis. Soluble p62 levels were significantly increased by 69 ± 21% in SCA3-CN relative to WT-CN (*p* = 0.03), and the ratio of insoluble to soluble p62 protein was more than doubled in SCA3-CN (*p* = 0.02) (Figure 7A and 7B). Thus, similar to SCA3-hESC, SCA3-CN exhibit increased p62 expression with some of it likely sequestered into insoluble ATXN3^+^ aggregates. Immunoblot analysis of the autophagy proteins Beclin1 and LC3 was also performed on unfractionated protein lysates from WT- and SCA3-CN. Beclin1 levels trended towards decreased levels but failed to reach statistical significance (*p* = 0.07), and LC3-II levels were unchanged in SCA3-CN relative to WT-CN (*p* = 0.72) (Figure 7C and 7D). Therefore, while SCA3-hESC and SCA3-CN both exhibit markers suggesting impaired autophagy, the observed differences suggest mutATXN3 may interact and interfere with autophagic pathways differently in undifferentiated hESC versus differentiated neurons.

**Figure 7.**
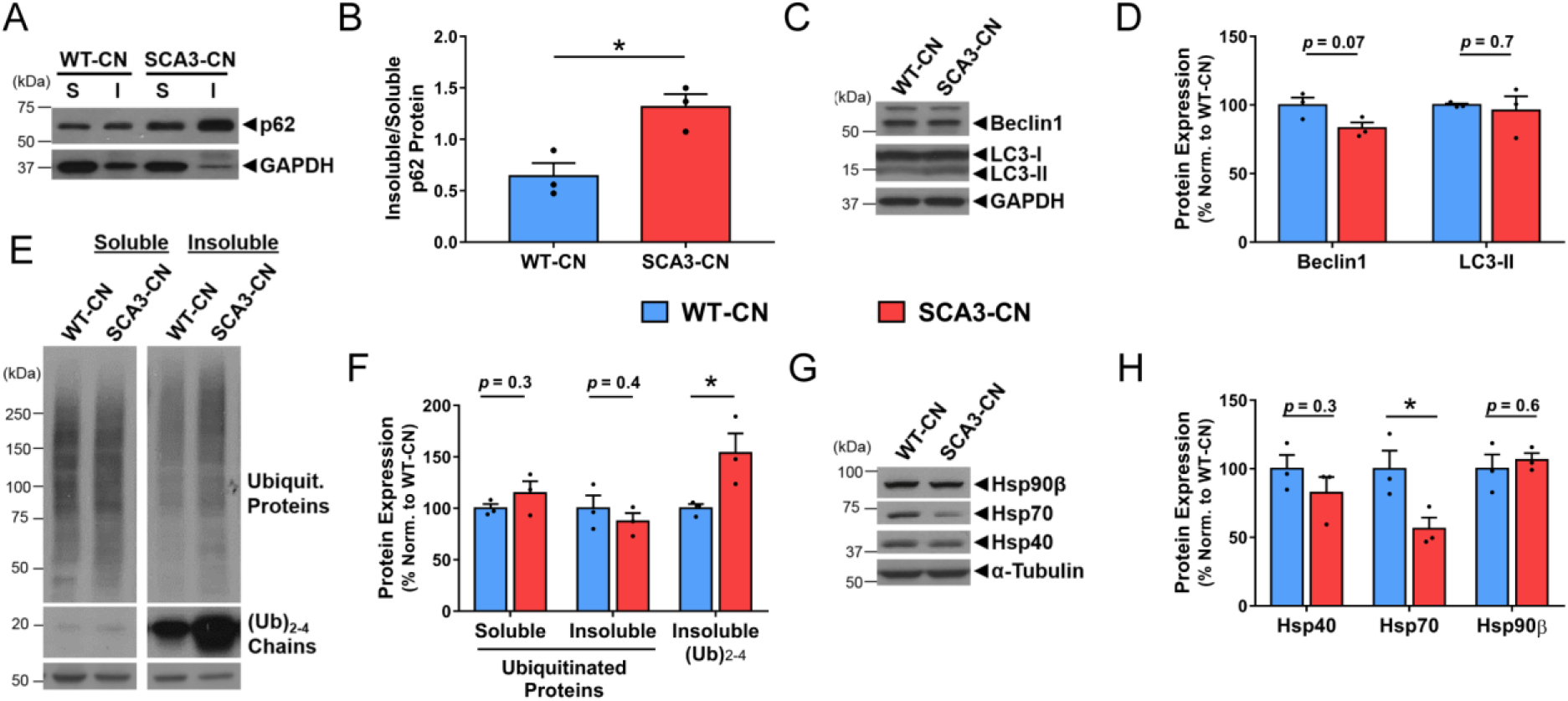
SCA3-hES derived Day 30 cortical neurons exhibit altered expression of key regulators of protein homeostasis. (A) Representative anti-p62 Western blot of soluble (S) and insoluble (I) protein fractions from WT-CN (blue) and SCA3-CN (red). (B) Ratio of insoluble/soluble p62 protein in WT-CN and SCA3-CN. (C) Representative anti-Beclin1 and anti-LC3B Western blots and (D) quantification in RIPA-lysed WT-CN and SCA3-CN. (E) Representative anti-ubiquitin Western blot of soluble and insoluble protein fractions from WT-CN and SCA3-CN. (F) Quantification of soluble ubiquitinated and insoluble ubiquitinated proteins, and insoluble small ubiquitin chains [(Ub)_2-4_] in WT-CN and SCA3-CN. (G) Representative anti-Hsp40, -Hsp70, and -Hsp90β Western blot and (H) quantification in RIPA-lysed WT-CN and SCA3-CN. Data (mean of 3 replicates ± SEM) are reported relative to mean WT-CN set to 100%. Unpaired two-tailed t test (\**p*<0.05). (Ubiquit. = ubiquitinated; GAPDH = glyceraldehyde-3-phosphate dehydrogenase).

Additional components of the protein homeostasis machinery were evaluated in SCA3-versus WT-CN. Anti-ubiquitin immunoblot analysis revealed no difference in total soluble or insoluble poly-ubiquitinated substrates in protein extracts from the two lines (Figure 7C and 7D). There was, however, a significant increase in the level of short ubiquitin chains [(Ub)_2-4_] partitioning in the insoluble fraction in SCA3-CN: a 54% ± 20% increase relative to WT-CN (*p* = 0.05) (Figure 7E and 7F). As in SCA3-hESC, Hsp70 levels were significantly decreased in unfractionated SCA3-CN relative to WT-CN (decreased by 44% ± 15%; *p* = 0.048) (Figure 7G and 7H). In contrast, Hsp40 and Hsp90β levels did not statistically differ between the two lines (*p* = 0.32 and *p* = 0.60, respectively) (Figure 7G and 7H). Overall, SCA3-CN exhibited significant changes in several markers of autophagy, ubiquitin-dependent proteasomal clearance, and protein folding. Several disease-specific changes in SCA3-CN mirrored those seen in undifferentiated SCA3-hESC while others did not, suggesting cell-type specific differences in the interaction of mutATXN3 with components of protein homeostasis.

## 4. Discussion

We present here a new SCA3 human embryonic stem cell line with the potential to improve our understanding of this complex, fatal neurodegenerative disease. Despite over 20 years of research since the identification of the disease mutation, limited understanding of SCA3 disease mechanisms and the absence of effective therapeutic interventions point to the need for improved humanized disease model systems^1^. Here we developed and characterized SCA3-hESC, a significant cell line partly because it is the only NIH-approved hESC line harboring a SCA mutation. More importantly, SCA3-hESC represents the first hPSC or cell model of any kind that displays disease-relevant molecular phenotypes, including robust aggregation of the endogenous human disease gene product, without the need for genetic or environmental manipulations^12,33–36^.

We observed multiple molecular phenotypes at the undifferentiated hESC stage, including the formation of p62-positive aggresomes containing the disease protein ATXN3 and altered expression of key regulators of protein homeostasis. These findings were unexpected as SCA3 typically presents later in life with selective neuronal vulnerability^1,4,5^. The presence of disease phenotypes in undifferentiated cells could point to early developmental changes that are as yet unappreciated in SCA3, and is reminiscent of recent findings in HD, another adult-onset polyQ disorder^55^. Cellular properties of pluripotent stem cells, notably their heavy reliance on autophagy-dependent protein clearance and rapid rate of division, may also provide insights into polyQ expansion-driven changes in ATXN3 function. Identification of disease protein aggregation and altered protein clearance pathways in SCA3-hESC are also notable since no similar disease-relevant phenotypes have been observed in other published SCA3 iPSC lines without application of cellular stressors^12^. The apparent lack of robust phenotypes in SCA3 iPSC lines could arise from the reprogramming process from a differentiated cell, which can cause unpredictable changes in epigenetic patterning, transcriptional regulation, pluripotent potential, and vulnerability to early senescence^56^. Regardless, our findings bolster support for the development of more SCA3 hESC lines in addition to patient-derived iPSC lines to improve our understanding of SCA3 pathogenesis.

SCA3-hESC presumably can express all ATXN3 isoforms generated from the endogenous human disease gene and, as demonstrated here, recapitulates aggregation and nuclear sequestration of mutATXN3. Thus, SCA3-hESC is well suited to assess compounds aimed at reducing mutATXN3 protein, preventing aggregation, or altering subcellular localization of mutATXN3. Here we performed a proof-of-concept assessment of a previously validated gene silencing therapy^20,21^; specifically, we demonstrated that ASO-mediated reduction of ATXN3 mitigates aggresome formation in SCA3-hESC. The ability to use tools such as ASOs to modify disease-relevant phenotypes supports the hypothesis that at least some of the observed molecular phenotypes are directly attributable to mutATXN3 expression, and could enable future mechanistic studies of disrupted pathways in SCA3. However, we cannot rule out the possibility that some of the observed molecular phenotypes in SCA3-hESC and differentiated cell types are due to cell line-specific variations not directly resulting from expression of mutATXN3. Future studies using this novel cell line would benefit from development of genetically corrected isogenic control lines from SCA3-hESC, such as those that have been developed from Huntington’s disease iPSC lines^57^. In addition, the SCA3 field would benefit from the creation of more SCA3 hESC lines, which may occur due to increased numbers of patients choosing in vitro fertilization with preimplantation genetic testing for monogenic and repeat expansion disorders.

We also observed the formation of high molecular weight ATXN3 aggregates throughout neuronal differentiation, including in derived SCA3-NPC and mature SCA3-CN. Though there is evidence for involvement of the frontal, parietal, temporal, occipital and limbic lobes in SCA3^58,59^, the degree of neuronal loss in these regions is significantly less severe than in brainstem and cerebellar nuclei^60^. Because cortical regions likely play less of a role in the progressive deterioration of motor control in SCA3 patients, differentiated forebrain cortical neurons may not be the best cellular model in which to study SCA3 disease processes. Dopaminergic neurons in the substantia nigra, however, are highly affected and exhibit significant degeneration in SCA3 human brains. Future studies could take advantage of the wealth of improved dopaminergic differentiation protocols^61^ to assess specific differences in pathology and neuronal vulnerability in disease-affected dopaminergic versus disease-spared cortical neuronal populations. In addition, SCA3 research to date has been neuron-centric, meaning that little is known about glial involvement in disease progression. Recent studies demonstrating significant transcriptional and protein expression changes in glial-enriched genes^62–64^and altered glial morphology and density in affected brain regions with disease progression^4,65^ point to the need for glial-focused investigations in SCA3. The pluripotent potential of SCA3-hESC in combination with the detection of native mutATXN3 aggregation demonstrated here, could prove invaluable for future studies of cell-type specific differences in ATXN3 function, aggregation propensity and vulnerability to aggregation-induced cellular insults.

## 5. Conclusions

Here we have presented a novel hESC line that expresses the endogenous SCA3 human disease protein. We have verified the pluripotent potential of the SCA3-hESC line, and that it can be directed to differentiated into mature, electrophysiologically active neurons. Most importantly, we have identified robust, well-established and mutATXN3-dependent pathogenic features of SCA3 human disease in undifferentiated stem cells and throughout neuronal differentiation. Based on these studies, we believe the SCA3-hESC line holds strong promise to advance our understanding of SCA3 disease mechanisms, enable future investigations into cell-type specific contributions to disease and improve preclinical development of SCA3 therapies.

## Acknowledgements

We thank Ionis Pharmaceuticals for generating and providing the anti-*ATXN3* ASO. L.R.M., G.D.S., and H.L.P. conceived and designed the study. L.K. and G.D.S. generated the SCA3-hESC line from the donated human embryo. L.R.M., L.K., D.D.B., R.D., D.L., M.C.C., and H.S.M. performed the experiments. L.R.M. analyzed the data. V.G.S., G.D.S., and H.L.P. provided supervision for experimentation and data analysis. L.R.M. wrote the manuscript with immense support provided through discussions with and supervision by G.D.S. and H.L.P. All authors read and approved of manuscript.

